# Mitochondria, mutations and sex: a new hypothesis for the evolution of sex based on mitochondrial mutational erosion

**DOI:** 10.1101/019125

**Authors:** Justin C. Havird, Matthew D. Hall, Damian K. Dowling

**Affiliations:** Dept. of Biological Sciences, Auburn University, Auburn, AL 36849, USA; Biology Dept., Colorado State University, Fort Collins, CO 80523, USA; School of Biological Sciences, Monash University, Victoria 3800, Australia

## Abstract

The evolution of sex in eukaryotes represents a paradox, given the “two-fold” fitness cost it incurs. We hypothesize that the mutational dynamics of the mitochondrial genome would have favoured the evolution of sexual reproduction. Mitochondrial DNA (mtDNA) exhibits a high mutation rate across most eukaryote taxa, and several lines of evidence suggest this high rate is an ancestral character. This seems inexplicable given mtDNA-encoded genes underlie the expression of life’s most salient functions, including energy conversion. We propose that negative metabolic effects linked to mitochondrial mutation accumulation would have invoked selection for sexual recombination between divergent host nuclear genomes in early eukaryote lineages. This would provide a mechanism by which recombinant host genotypes could be rapidly shuffled and screened for the presence of compensatory modifiers that offset mtDNA-induced harm. Under this hypothesis, recombination provides the genetic variation necessary for compensatory nuclear coadaptation to keep pace with mitochondrial mutation accumulation.

## Introduction: The evolution of sex and the evolution of mitochondria are intrinsically linked

Widespread sexual reproduction among eukaryotes is puzzling from an evolutionary standpoint, because sex carries a “two-fold” fitness cost, and must therefore offer a benefit that surpasses this cost [1]. Several plausible models of the evolution of sex have been proposed, based on the benefits of recombination with sex, including facilitating the purging of deleterious mutations from the nuclear genome [2] or adaptation to ecological hazards such as parasites or environmental fluctuations [3]. However, while experimental and theoretical support exists for each of these classes of models, they arguably remain limited in their capacity to explain certain fundamental questions: why have all eukaryotes, but no prokaryotes, evolved life-history strategies hinged on sexual reproduction; and why has obligate sexual reproduction become so prevalent amongst certain eukaryotes (e.g., most metazoans), while other taxa (e.g., plants and certain metazoan lineages) exhibit more flexible reproductive systems based on facultative sex interspersed with clonal reproduction?

Eukaryotes are bound by two basal features. They all have – or have had during their evolutionary histories – energy-converting organelles called mitochondria [4]; and they all have – or have had – the ability to reproduce sexually [5]. We propose that the origins of this ancient association can be traced to the mutational properties of the mitochondrion’s own (mt)DNA. The mitochondria are integral to eukaryote evolution, and the ancient endosymbiosis that led to the mitochondrion provided the eukaryotes with a highly efficient form of energy conversion, presumably catalyzing the evolution of complex life [6]. However, mitochondria retain a small genome, which is destined to accumulate mutations via Muller’s Ratchet as a consequence of the genome’s evolutionary constraints and high mtDNA mutation rate across the eukaryote phylogeny [7,8] (with few exceptions, such as the derived slow mutation rate of many land plants). Furthermore, across the eukaryote domain, de novo mutations in the mtDNA exhibit a higher fixation probability than those in the nuclear DNA [9]. This leads to a striking paradox – a genome that wields a salient hand in maintaining the integrity of complex life, is prone to perpetual mutational erosion across large branches of the eukaryote phylogeny.

Here, we outline a new hypothesis that can reconcile this paradox. We propose that mitochondrial genomic mutation accumulation (hereafter termed “mito-mutation accumulation”) is likely to have represented one of the original drivers of the evolution of sexual reproduction in eukaryotes. We contend that during the early evolution of eukaryotes, mito-mutation accumulation would have placed strong selection on the host genome for compensatory modifier mutations to offset the negative metabolic effects linked to deleterious mtDNA mutations. Recombination between distinct host ‘nuclear’ genotypes, achievable via sexual reproduction, would facilitate such a compensatory response, by accelerating the rate by which alleles at host nuclear loci could be shuffled and screened for compensatory function.

Below, we outline our case, which is built on a strong base of experimental studies that have elucidated how compensatory mitochondrial-nuclear (mito-nuclear) coevolution is key to maintaining organismal viability. We highlight evidence for nuclear compensatory adaptation to pathogenic mtDNA mutations, and discuss how mito-nuclear coevolution is greatly facilitated by recombination in the nuclear genome. We then identify the key assumptions on which our hypothesis rests, and present supporting evidence for each assumption. We discuss the hypothesis in the context of other evolutionary theories that have previously drawn links between the mitochondria and sex in eukaryotes. Finally, we conclude by noting that the power of our hypothesis lies in its testability; it provides a series of predictions amenable to experimental enquiry, whose answers could provide compelling support for the contention that mito-mutation accumulation was a key driver of the evolution of sex in eukaryotes.

## Experimental support for the role of compensatory nuclear adaptation to mito-mutation accumulation

Preserving the fine-scale interactions between proteins encoded by both mitochondrial and nuclear genomes is critical for maintaining oxidative phosphorylation (OXPHOS) and meeting the energy needs of the contemporary eukaryotic cell. Paradoxically, the mtDNA is prone to perpetually accumulate deleterious mutations [10], which could have fatal consequences for organismal function in the absence of a compensatory nuclear response [11-16]. Indeed, it is now well known that mutations in the mtDNA sequence are often tied to the onset of metabolic diseases, early ageing, and infertility [17,18]. Moreover, an increased penetrance of such ailments, and a general reduction in fitness, has been observed upon artificially disrupting coevolved combinations of mito-nuclear genotypes, by expressing mtDNA haplotypes alongside evolutionary novel nuclear backgrounds [19,20]. In model systems like *Mus* and *Drosophila*, such mito-nuclear “mismatches” have resulted in reduced metabolic functioning and lower fitness [16,21-23]. These results strongly suggest that, within any given population, mitochondrial and nuclear gene combinations are co-evolutionarily “matched” to one another. It follows that mito-nuclear coevolution may contribute to driving reproductive isolation between incipient populations, and that this might ultimately lead to speciation [12,24].

Other support for the role of mito-nuclear coevolution in maintaining organismal viability comes from studies reporting elevated substitution rates in nuclear-encoded genes that interact with mtDNA-encoded products, relative to their nuclear counterparts that interact with other nuclear-encoded genes [25-27]. These findings are important because they indicate a key role for positive selection in shaping the genetic architecture of nuclear-encoded genes involved in mitochondrial function in order to compensate for mtDNA mutations. Given the products of these nuclear genes are entwined in tightly regulated mito-nuclear interactions, this suggests that these nuclear genes are responding quickly and efficiently to mito-mutation accumulation, via counter-adaptations of compensatory function to ensure the integrity of metabolic function [25,28].

The empirical evidence outlined above suggests that nuclear compensatory adaptations have commonly evolved to mitigate the effects of mito-mutation accumulation and prevent mtDNA-mediated Muller’s Rachet effects from leading to widespread lineage extinctions across the eukaryote phylogeny. Recombination in the host genome would have greatly facilitated this compensatory response, by shuffling and generating unique nuclear allelic combinations every generation, providing the genetic variation required for selection to screen for nuclear adaptations that offset mitochondrial mutational erosion (Fig. 1). In early eukaryotes, while horizontal gene transfer from other host nuclear lineages might have partially alleviated the effects of mito-mutation accumulation, we contend that a more predictable and rapid mechanism of gene mixing would have been required to cope with the perpetual mutational pressure imposed by the mitochondrial genome during this time (Fig. 1). Recombination via sex could provide such a mechanism, enabling compensatory nuclear alleles to spread quickly into new genetic backgrounds, offsetting Hill-Robertson effects [29], and ensuring the modifiers were not placed on nuclear genomic backgrounds that are largely deleterious in performance and hence purged under background selection.

**Figure 1.**
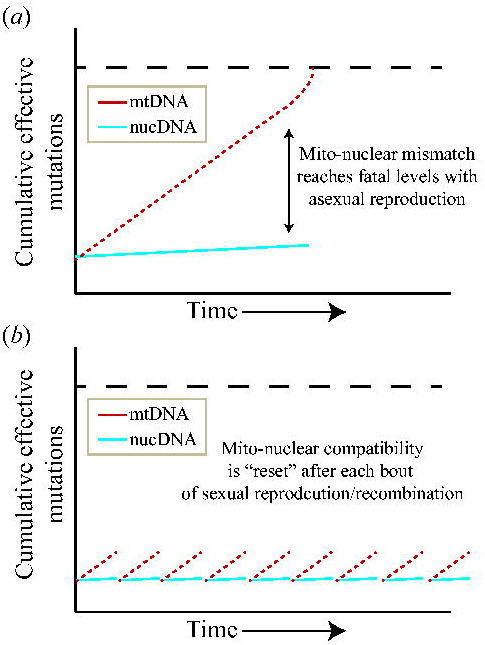
Mutational meltdown in the mitochondrial DNA (mtDNA) is offset by compensatory changes in the nuclear DNA, facilitated by recombination associated with sexual reproduction, providing a mechanism for the origin and maintenance of sexual reproduction. **A:** During asexual reproduction, mito-nuclear mismatch increases over generations as mtDNA accumulates mutations faster than nucDNA. Here, we envisage the mtDNA mutation rate might ultimately increase exponentially in the absence of a nuclear compensatory response, once mito-nuclear mismatch reaches a threshold. This is because the accumulation of mtDNA mutations per se might act as a catalyzing force for further mutation accumulation (if for instance there is a positive association between mtDNA mutation numbers and reactive oxygen species production rates). **B:** With sexual reproduction, organismal viability is restored in the population by selection for compensatory genotypes in the nucDNA, facilitated by recombination. Red/dotted lines represent mtDNA, blue/solid lines represent nucDNA, and dashed lines represent a fatal upper threshold for mito-nuclear mismatch. “Cumulative effective mutations” is shown on the y-axis, and denotes the effects of these mutations on the phenotype. Even though the absolute number of mutations in the mitochondrial genome (and nuclear genome) remains the same in panel (a) and (b), in (b) the mutations are matched by nuclear compensatory adaptations in the recombined nuclear genome.

## Key assumptions

Our hypothesis centres on two assumptions, each of which are plausible and which have been made previously by evolutionary biologists. We outline the substantiating evidence for each below. Furthermore, we note that while there are exceptions to each of these assumptions, it is these very exceptions that provide excellent opportunities on which our hypothesis can be tested (see Predictions).

### Assumption 1: The high mutation rate of the mtDNA is an ancestral condition

We assume that the high mutation rate of mtDNA, which is a hallmark of the streamlined mitochondrial genomes of metazoans [30], is a character that was shared by ancestral mitochondria during early eukaryote evolution and the progression of endosymbiosis [31]. Six lines of evidence support this assumption. First, recent studies have identified unicellular and early-diverging eukaryotic lineages (e.g., haptophyte and stramenopile algae) that similarly show elevated mtDNA mutation rates, suggesting that across eukaryotes as a whole there is a propensity for mtDNA evolutionary rates to outstrip those in the nucleus [7,8,32]. Fungi and yeast species also conform to these high mtDNA mutation rates relative to nuclear rates [33-35], indicating that elevated mtDNA mutation is not restricted to animals. Together, these data suggest that a high mutation rate is likely to have been shared by the eukaryotic ancestor, and that the low mtDNA mutation rate of many land plants is a derived condition. Indeed, studies have emerged that demonstrate low mtDNA mutation rates are not systematic of all land plants, and many such species show incredible variation in the mtDNA mutation rate, including some of the highest ever documented rates recorded in eukaryotes [36,37].

Second, other recent cellular endosymbionts likewise exhibit elevated mutation rates compared with the genomes of their hosts, suggesting that during endosymbiosis the ancestral mitochondria would have also experienced increased mutation rates [38,39]. Third, rates of substitution appear to be faster in bacteria, which gave rise to the mitochondrial genome, than in archaea, which gave rise to a majority of the nuclear genome [40]. Fourth, the well-documented presence of a “long-branch” leading from the bacterial ancestor to the mtDNA of modern eukaryotes (including plants) suggests a rapid increase in mutation rate in the mtDNA of early eukaryotes following endosymbiosis [41], further supporting the conclusion that the slow-evolving mtDNA of plants is a derived characteristic. Fifth, we note that the mtDNA resides within the mitochondria – a highly mutagenic site that is the major source of reactive oxygen species in the cell due to its redox activity. Reactive oxygen species production also increases during ageing and is correlated with compromised mitochondrial function [42]. Finally, given that mtDNA replicates more often than the nuclear DNA per cell cycle [43], replication errors should be more prevalent in the mtDNA, leading to an increased mutation rate [18] across the eukaryote phylogeny.

### Assumption 2: Fundamental evolutionary constraints limit the mtDNA itself from evolving bi-parental transmission

Across eukaryotes, uniparental inheritance of the mitochondria remains a ubiquitous pattern, with only a few exceptions [44]. Our hypothesis assumes the mtDNA was under strong evolutionary constraints to avoid bi-parental transmission, as its host transitioned to sexual reproduction with two sexes. We acknowledge that there is considerable flexibility in mtDNA recombination rates across eukaryotes, particularly in land plants and in fungi. However, intra-individual heteroplasmy of divergent mtDNA molecules in plants is rarely reported to be caused by paternal leakage (as a consequence of sexual reproduction), and uniparental transmission of mtDNA remains the typical pattern in in these lineages [45]. This is similarly the case for fungi [46], which show heteroplasmy in early life stages, but revert to homoplasmy as they develop [45]. Even in these taxa in which recombination of mtDNA has been recorded [45], they maintain uniparental inheritance of the mtDNA, which would presumably reduce the scope by which recombination could act to purge poorly performing mtDNA molecules. Genetically effective recombination between mtDNA molecules requires bi-parental transmission. In the absence of biparental transmission, any two mtDNA molecules would share near-identical sequences, and so too would the products of their recombination [43].

The assumption that fundamental evolutionary constraints prevented the evolution of biparental mitochondria transmission has received robust support from population genetic theory [47-50], as well as empirical work [51,52]. Population genetic models have demonstrated that biparental transmission would lead to divergent mtDNA molecules competing within the same host, and promote selection for selfish molecules that replicate faster at the expense of host metabolic efficiency [47-50]. Exploring these conditions, experimental studies in yeast and mice have shown that heteroplasmy leads to competition between divergent mtDNA genomes and a breakdown in fundamental metabolic functions such as respiration, nutrient intake, and even cognitive impairment [51,52]. Additionally, maintaining uniparental inheritance allows for mtDNA to be sequestered in a metabolically quiescent germ-line, which allows resulting offspring to avoid inheriting a damaged mtDNA genome associated with a metabolically active cell (e.g., sperm cells) [44,53,54].

Finally, while prokaryotes rely on horizontal gene exchange as a facilitator of adaptation, we assume this level of gene-mixing would not have sufficed the evolving eukaryote. Transitioning to life as a highly complex cell, and harnessing the efficient energy conversion provided by the mitochondria, is predicted to have led to a rapid expansion in nuclear genome size [6]. As the nuclear genome expanded, however, horizontal gene transfer would have affected a smaller and smaller fraction of the nuclear genome, diminishing the capacity of this process to reliably provide the raw genetic variation needed to screen for compensatory nuclear modifiers. Indeed, only one asexual eukaryotic lineage, the bdelloid rotifers, is believed to experience a large amount of horizontal gene transfer as a possible substitute for sex [55]. Yet rotifers also have high rates of gene conversion and expanded numbers of genes for oxidative resistance [56], suggesting that the lack of recombination in eukaryotes is exceedingly rare because multiple mechanisms, and a very specific genetic architecture, is required to supplement horizontal gene transfer.

## Previous links between mitochondria and the evolution of sex

A link between mitochondria and sex [6,49,50,57-59] has been suggested in previous studies, but these have typically assumed gamete fusion (i.e, sex) was already in place at the time of acquisition of the mitochondrion. Here, the evolution of two different sexes – one that transmits the mitochondria (the female), and one that does not (the male) – mitigates the selfish conflict between divergent mtDNA molecules of different parents, by maintaining uniparental mtDNA transmission [49,50,58,59] (see Assumption 2 above). None of these studies envisioned that acquisition of the mitochondrion, in early eukaryote evolution, was a driving force behind the evolution of sex *per se*. Only one other viewpoint has previously proposed a link between recombination and mitochondria, albeit with a different process mediating the evolution of sex [57]. Lane [57] suggested the early mitochondrion in the evolving eukaryote would provide an unending source of foreign DNA that would have contaminated the nuclear genome, and this contamination of foreign DNA would have imposed selection for recombination between host genotypes to preserve nuclear chromosomes with beneficial mutations, while purging deleterious ones. This view is supported by evidence that introns in the nuclear genome are of mitochondrial origin [60]. Lane and Martin [6,61] also hypothesized that the expansion of nuclear genome size, facilitated by the mitochondrion, would have favoured a shift from unreliable lateral gene transfer to the more robust sexual recombination [6].

Our hypothesis differs from those of predecessors by proposing that the selection for sex and nuclear recombination in eukaryotes came from mutational meltdown within the mitochondrial genome itself. By recognizing that mitochondrial mutations are particularly consequential because they are more often fixed than nuclear mutations [9], and because they affect one of eukaryote life’s core functions – energy conversion, we extend the Fisher-Muller model of recombination to include the fixation of beneficial modifiers that together act to offset the effects of mito-mutation accumulation [62]. Moreover, we note that parasite-mediated theories of sexual reproduction [3] have clear parallels to our hypothesis, given that the ancestral mitochondria can reasonably be envisaged as endosymbiotic parasites. As with any complex inter-species interaction [63], early eukaryotic-mitochondrial symbiosis may have been marked by pervasive inter-genomic conflict, signatures of which are still manifest in contemporary eukaryote lineages (e.g. cytoplasmic male sterility in plants [64]). Thus, in addition to the rapid accumulation of mito-mutations, early eukaryotic-mitochondrial conflict might conceivably have selected for mutations that benefited the mitochondria, at the expense of host fitness, further promoting the evolution of recombination in the host nuclear genome to enable an efficient response to the evolving mtDNA.

Although formation of chimeric OXPHOS complexes between mitochondrial and nuclear proteins would have greatly spurred the need for exquisite mito-nuclear compatibility [15,16,65], we believe that the evolutionary transition to sexual reproduction in order to maintain mito-nuclear compatibility is very likely to have pre-dated the origin of OXHPOS complexes. Parts of the metabolic machinery of eukaryotes would have required intimate inter-genomic coordination to achieve mutual endosymbiosis, long before OXPHOS-encoding genes translocated over to the nuclear genome. For example, the nuclear-encoded genes involved in importing raw materials from outside of the host and into the mitochondria, would presumably have needed to co-evolve with mtDNA-encoded genes involved in converting and utilizing such materials. Indeed, we contend that it was the transition to host nuclear recombination and sex that would have provided the large selective advantage for translocated mtDNA genes to move over *en masse* and remain in the nucleus (Fig. 2), resulting in the greatly streamlined mito-genomes of eukaryotes compared to their bacterial ancestors.

**Figure 2.**
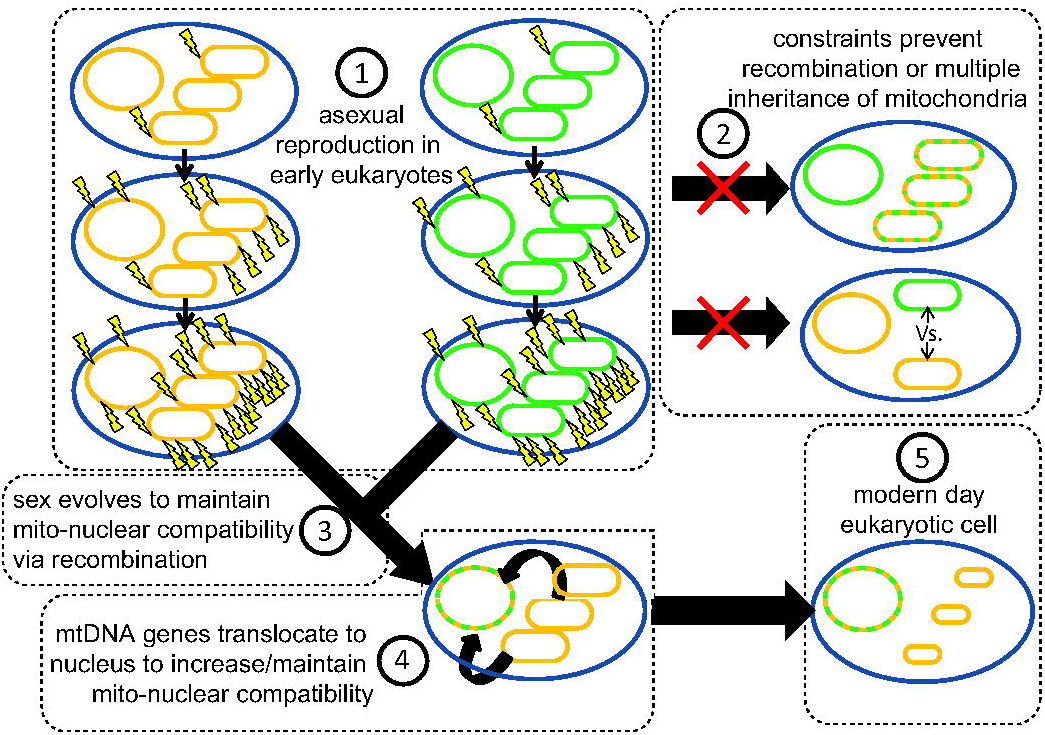
A model for early eukaryotic evolution. Under our hypothesis, the evolution of recombination via sexual reproduction stems from the need to shuffle nuclear genes, as an inevitable consequence of having mitochondria. We envisage that this was driven by high mtDNA mutation rates in the early endosymbiotic bacterium that evolved into the mitochondrion. Following acquisition of the bacterial endosymbiont, the early eukaryote would have transmitted its endosymbiont vertically (number 1), without recombination, leading to mutational erosion in the mtDNA. Numbers indicate the general temporal order of other events, including the evolution of sex, based on our hypothesis. Lightning bolts denote DNA mutations and different colors represent DNA from different lineages. Circles denote host nucleus and associated genome, and the eclipses denote mitochondria and their mtDNA.

## Testable predictions and future experiments

There are several testable predictions that stem from our hypothesis. First, we predict that eukaryote taxa with lower mtDNA mutation rates should exhibit lower propensities for sexual reproduction. Although mtDNA mutates quickly in most animal lineages, some (e.g., Cnidaria) have slowly-evolving mtDNA [66], and Angiosperms in general also have slowly-evolving mtDNA [8]. Under our hypothesis, the magnitude of mtDNA-mediated selection for compensatory nuclear modifiers should be lower in these lineages. Cursory evidence suggests obligate sexual reproduction and outcrossing are relatively rare in both lineages [67,68], supporting the prediction that lineages with low mtDNA mutation rates should not have to rely as heavily on sexual reproduction to offset mtDNA-induced harm. We would expect this correlation to extend across diverse eukaryotic lineages.

A second prediction can be derived based on the rate of paternal leakage of mtDNA, and associated recombination, across eukaryotes. Although mtDNA is overwhelmingly transmitted through the maternal lineage in eukaryotes, rare cases have been documented of paternal mtDNA also being transmitted, suggesting recombination between mtDNA genomes to prevent mito-mutation accumulation might be possible over evolutionary timescales in some lineages [44]. Those lineages in which leakage of mtDNA from the paternal parent is prevalent should be able to resort to this mechanism in part to offset their mitochondrial mutation loads, as suggested previously [44,69]. Based on our hypothesis, it follows that species with appreciable levels of parental mtDNA leakage should also be less likely to exhibit modes of obligate sexual reproduction.

Third, experimental disruption of coevolved mito-nuclear genotypes in facultatively sexual species should induce elevated rates of sexual reproduction in these species. Experimental mismatching of coevolved mito-nuclear genotypes, achieved by pairing the prevailing mtDNA haplotype of a population or species alongside an evolutionary novel nuclear genome sourced from a separate population/species, has been shown to decrease organismal fitness and modify patterns of gene expression [16,19-23,70,71], but has never previously been harnessed to study the propensity for sexual reproduction. Here, the assumption is that the creation of mito-nuclear mismatches will promote individuals to resort to sex to harness the benefits of recombination between divergent host nuclear genotypes, to optimize selection for counter-adaptations that restore mito-nuclear compatibility.

Finally, under the assumption that the nuclear-encoded genes that directly interact with those encoded by the mtDNA are more likely to host compensatory mutations, these nuclear-encoded genes are predicted to exhibit elevated recombination rates relative to other nuclear genes that are not involved in mito-nuclear interactions. As we noted above, there is already evidence that nuclear gene products that interact with mtDNA-encoded products evolve more quickly based on studies of primates, copepods and plants, supporting this prediction [25-27]. Recent technological developments enabling deep-sequencing to identify recombination “hot spots” across the nuclear genome [72] could be used to directly gauge recombination levels in mitochondrial interacting genes relative to those that do not interact with the mitochondria.

## Conclusions

We have presented a new hypothesis for the evolution and maintenance of sexual reproduction. This hypothesis draws on a robust foundation of empirical evidence that has substantiated the evolutionary consequences of mito-mutation accumulation, and the role of mito-nuclear interactions in population coevolutionary processes. Backed by this evidence, we note that the mitochondrial genome – a universal feature of all organisms equipped for sexual reproduction – is destined to accumulate mutations. We posit that this would have invoked selection in the ancestral eukaryote for a mechanism that would facilitate the rapid screening of new combinations of nuclear genotype, for those exhibiting compensatory function to offset this mitochondrially-induced harm. We propose that this selection pressure was a catalyst for the evolution of sexual reproduction, since sex, when combined with recombination, would provide the abundant novel genetic variation necessary for selection to evaluate genotypes of compensatory effect. The power of our hypothesis lies in its testability. The predictions that we have outlined are readily amenable to scientific testing. This can be achieved through experimental studies in modern eukaryote lineages that directly evaluate whether mito-nuclear genomic mismatches can be alleviated and restored via sexual recombination, as well as by correlative studies examining mutational characteristics of mitochondrial and nuclear genomes. Our hypothesis, which links mtDNA mutation rates and mito-nuclear conflict to the propensity for sex, offers an empirically tractable solution to an enduring problem in biology.

## Acknowledgements

We thank Daniel Sloan, Rachel Muller and their lab groups at CSU and the Mitonuclear Ecology discussion class at Auburn University for stimulating discussion of this work. DKD/MDH were funded by the Australian Research Council. The authors declare no conflicts of interest.

